# Rhamnogalacturonan-II dimerization deficiency impairs the coordination between growth and adhesion maintenance in plants

**DOI:** 10.1101/2024.11.26.625362

**Authors:** Abu Imran Baba, Lucija Lisica, Asal Atakhani, Bibek Aryal, Léa Bogdziewiez, Özer Erguvan, Adrien Heymans, Pawan Kumar Jewaria, Richard S. Smith, Rishikesh P. Bhalerao, Stéphane Verger

## Abstract

Cell adhesion is a fundamental feature of multicellular organisms. In plants, cell adhesion is mediated by the cell wall, but the control and maintenance of cell adhesion during growth and development remains poorly understood^1^. Here we uncover the role of a component of the cell wall, rhamnogalacturonan-II (RG-II) and its capacity to crosslink in the presence of Boron^2^, as a key regulator of plant cell adhesion maintenance. We show that RG-II dimerization deficiency leads to cell adhesion defects. Importantly, the analysis of *mur1* mutants with RG-II dimerization defects uncovers a cell adhesion pathway that is distinct from that identified by the analysis of pectin deficient mutants^3^. We found that mutations in two cell wall integrity sensors, RESISTANCE TO FUSARIUM OXYSPORUM 1 and RECEPTOR-LIKE PROTEIN 44, as well as supplementation with the hormone brassinosteroid can partially rescue the adhesion defects associated with RG-II dimerization deficiency. We also show that adhesion defects associated with RG-II dimerization deficiency are related to increased epidermal tension as well as decreased homogalacturonan levels in the cell wall, which can also be rescued by supplementation with brassinosteroid. Overall, we propose that RG-II dimerization defects alter cell adhesion directly (reduced crosslinks) but also indirectly through cell wall integrity sensing, brassinosteroid signalling, cell wall remodelling and cell layer growth coordination. Thus, our results uncover the involvement of cell wall integrity sensors and hormonal signalling in the coordination between growth and adhesion maintenance in plants, which is a key feature for complex multicellularity.

**Highlights:**

- RG-II dimerization is required for cell-cell adhesion in plants.
- Cell wall integrity sensors and brassinosteroid signalling mediate cell adhesion downstream of RG-II dimerization.
- Cell detachments due to defective RG-II dimerization are caused by weakened middle lamella and higher tissue tension.

## Result and discussion

Cell-cell adhesion in plants is often thought of as a passive outcome of cell wall synthesis. Adjacent plant cells adhere to each other via their cell wall and the middle lamella, a thin pectin layer enriched in homogalacturonan (HG), that is believed to play an adhesive role at the cell-cell interface^1,4–7^. This was supported by the characterization of mutants deficient in HG synthesis such as *quasimodo1* and *quasimodo2* that show cell adhesion defects^8–10^. Nevertheless, a suppressor screen on the *quasimodo* mutants identified that a mutation in *ESMERALDA1* can largely rescue *quasimodo* mutants’ adhesion defects without restoring the pectin content in the cell wall^3^. Along with subsequent work^11,12^, this suggested that adhesion maintenance in plant was in fact likely regulated by cell wall integrity sensing mechanism^1^, yet their exact nature and the molecular components involved have not been identified^13^. Rhamnogalacturonan-II (RG-II, another component of the pectin network) and its capacity to crosslink in the presence of boron has also been proposed to play a role in plant cell adhesion^14–16^, and could thus reveal a new pathway regulating adhesion in plants, but its actual contribution remains to be characterized. To test the contribution of RG-II dimerization for cell-cell adhesion in plants, we first studied the *murus1 (mur1)* mutant, which was previously shown to be deficient in pectin RG-II dimerization^17^. We grew wild type and mutant seedlings *in vitro* and in the dark to promote fast and extensive elongation of the hypocotyl. Using 3D confocal microscopy and cell wall staining with propidium iodide, we observed that two loss-of-function alleles, namely *mur1-1* and *mur1-2*, display cell adhesion defects (Figures 1 and S1). To reveal and quantify the extent of this adhesion defect on whole hypocotyls, we made use of ruthenium red staining^13^ and developed a workflow for high-throughput imaging and analysis of staining intensity and hypocotyl length. Our quantitative analysis confirmed a widespread staining (and thus defective epidermis) as well as shorter hypocotyl for *mur1-*2 compared to Col-0 (Figures 1 and S1). Along with our cellular-level confocal observations and similar observations in *mur1-1*, our results confirm that *mur1* mutants indeed display cell-cell adhesion defects.

**FIGURE 1.**
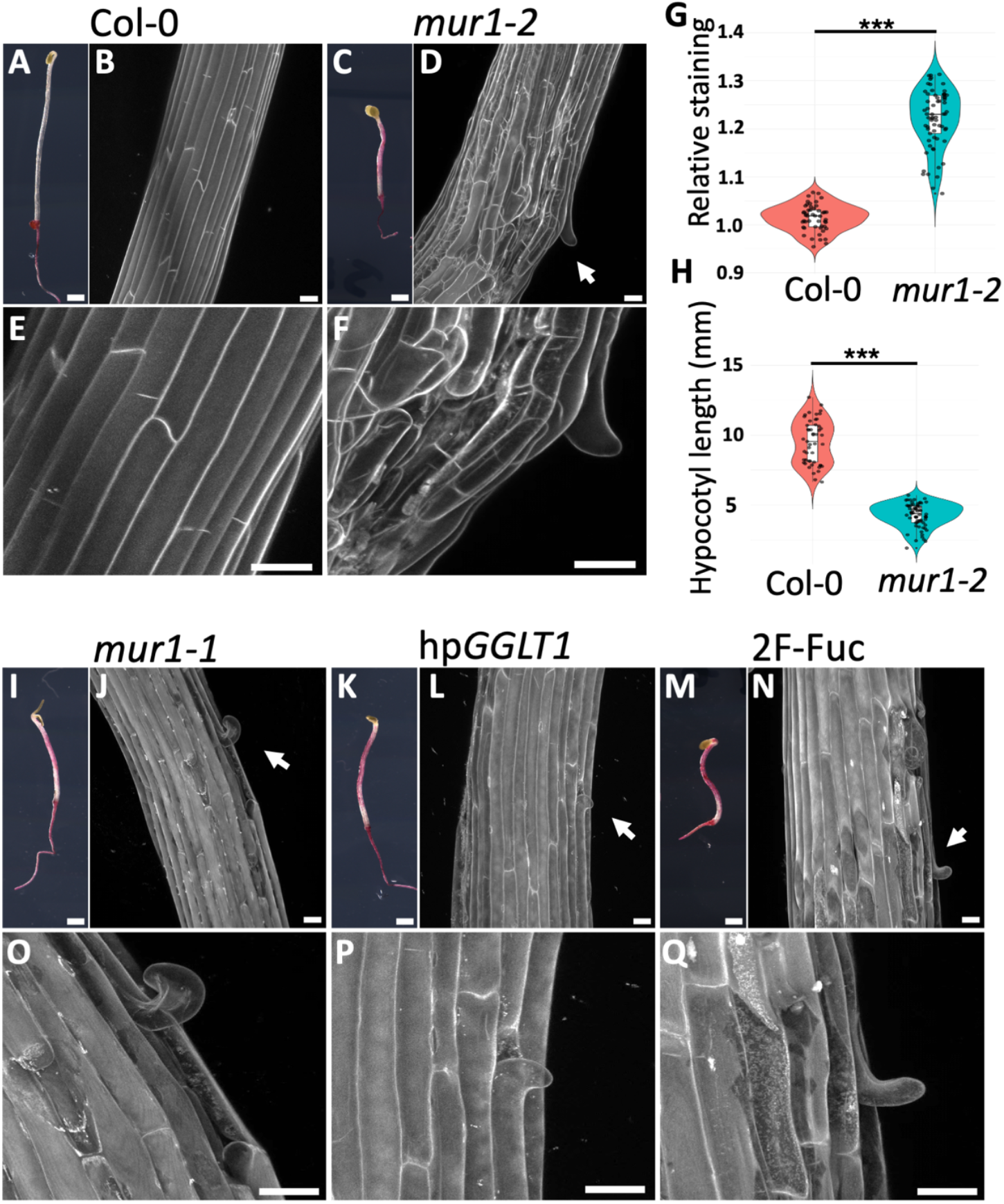
RG-II dimerization is crucial for cellular and supracellular integrity. (A, C, I, K and M) Representative ruthenium red staining images from 4-days-old dark-grown hypocotyls for Col-0 (A), *mur1-2* (C), *mur1-1* (I), hp*GGLT1* (Line #1) (K) and 15 µm 2F-Fuc treatment (M). (B, D, E, F, J, L, N, O, P and Q) Representative max intensity Z-projections of confocal stacks from 4-days-old dark-grown hypocotyl stained with propidium iodide from Col-0 (B and E), *mur1-2* (D and F), *mur1-1* (J and O), hp*GGLT1* (Line #1) (L and P) and 2F-Fuc (N and Q). Images in (E, F, O, P and Q) are magnifications highlighting the cell adhesion defects pointed by a white arrow in (B, D, J, L and N). (G) Quantification of the relative ruthenium red staining intensity and (H) measurement of hypocotyl length for Col-0 and *mur1-2*. Violin plot and boxplot summarize three biological replicates and asterisks indicate statistically significant difference between Col-0 and *mur1-2* determined by two-tailed Student’s t test, where ***P < 0.0005. [Scale bars, 30 µm in confocal images and 900 µm in Ruthenium red staining images.]. See Related content in figure S1 and S2.

Because *MUR1* encodes a GDP-mannose-4,6-dehydratase that catalyses the first step in *de novo* biosynthesis of GDP-I-Fucose^18^, we next tested if the phenotype observed was specific to an RG-II fucosylation defect or may be attributable to another fucosylated substrate. Known mutants for fucosyltransferases of xyloglucan^19^, arabinogalactan protein fucosylation^20^, protein α1-3 fucosylation^21^ and protein o-fucosylation^22,23^ did not show any visible adhesion defects (Figure S1). Yet treatment with the fucosyltransferase inhibitor 2-Fluoro-L-Fucose (2F-Fuc)^24,25^ mimicked the *mur1* mutants’ phenotype (Figure 1 M, N and Q). This suggest that one or several yet unidentified fucosyltransferases (as is the case for the RG-II specific fucosyltransferases) are involved in the adhesion phenotype observed. We were also able to largely rescue *mur1-2* adhesion phenotype by supplementation with borate (Figure S2), suggesting that the phenotype we observe in *mur1* can be rescued by the restoration of a normal level of boron-mediated RG-II dimer^17^. Finally, we also observed clear adhesion defects in two hairpin silencing lines (hp*GGLT1*) ^26^ for GOLGI GDP-l-GALACTOSE TRANSPORTER1 (GGLT1) that were shown to have RG-II specific galactosylation defects leading to impairments in RG-II dimerization^26^ (Figures 1 K, L, P and S1 V and W). All together our results strongly suggest that a deficiency in RG-II dimerization is sufficient to induce a loss of cell adhesion, indicating an important role for RG-II dimerization in cell-cell adhesion.

We next tested if adhesion defects in *mur1* can be attributed to the same pathway as those observed in the *qua* mutants. Interestingly, the double mutant *mur1-2 qua2-1* shows a strongly increased cell detachment phenotype and shorter hypocotyl when compared to *mur1-2* and *qua2-1* alone (Figures 2 and S3), suggesting independent pathways. We also tested boron supplementation on *qua2-1* which did not rescue *qua2-1* phenotype (Figure S2). Similarly, the *mur1-2 esmd1-1* double mutant did not show any rescue of *mur1-2* adhesion phenotype, as opposed to the rescue observed in *qua2-1 esmd1-1* (Figure 2 and S3). Overall, our results strongly suggest that *mur1* (and RG-II dimerization deficiency) affect adhesion through a pathway distinct from that mediated by homogalacturonans deficiency, characterised by analysis of *qua2 and esmd1*.

**Figure 2:**
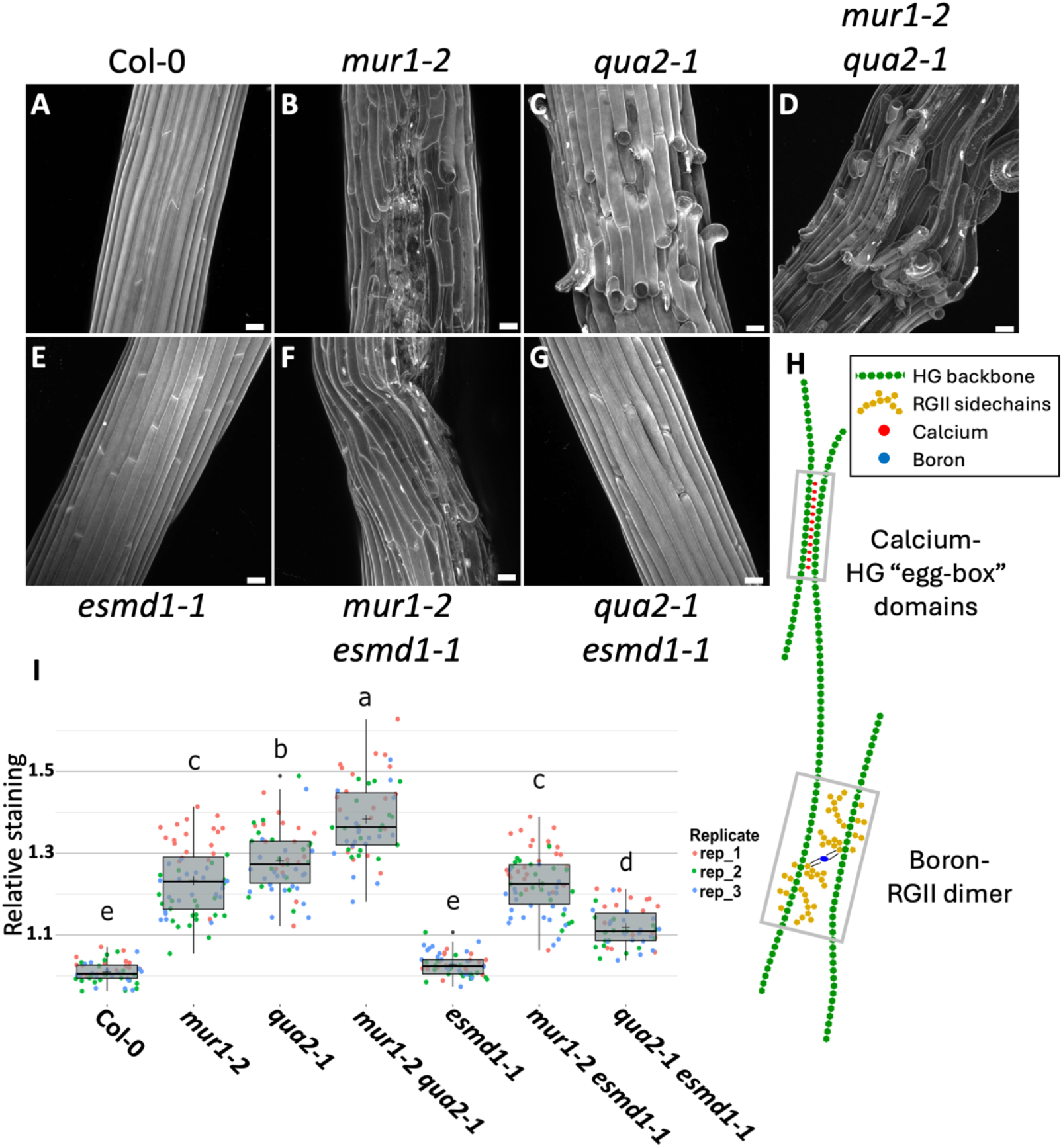
Homogalacturonan deficiency and rhamnogalacturonan-II dimerization defects affect adhesion through largely separate pathways. (A-G) Representative max intensity Z-projections of confocal stacks from 4-days-old dark-grown hypocotyl stained with propidium iodide from Col-0 (A), *mur1-2* (B), *qua2-1* (C), *mur1-2 qua2-1* (D), *esmd1-1* (E), *mur1-2 esmd1-1* (F) and *qua2-1 esmd1-1* (G). (H) Schematic representation of Boron-Rhamnogalacturonan-II (RG-II) dimers and Calcium-Homogalacturonan (HG) “egg-box” crosslinking domains along the HG backbone. (I) Quantification of the relative ruthenium red staining intensity (see representative images and length quantification in figure S3). Boxplots summarize three biological replicates, as represented by dots of different colors, and letters describe the statistically significant differences between populations determined by one-way ANOVA followed by Tukey’s HSD test (P < 0.05).[Scale bars, 30µm in images (A-G).] See related content in figure S2 and S3.

We next hypothesized that part of *mur1* adhesion phenotype may stem from secondary responses induced by cell wall integrity defects linked with reduced RG-II dimerization. We took advantage of the fucosyltransferase inhibitor 2F-fuc that we demonstrated to phenocopy *mur1* adhesion phenotype (Figure 1 M,N and Q) to screen a set of mutants for receptors previously proposed to act as cell wall integrity sensors^27^. We found that a mutant for RESISTANCE TO FUSARIUM OXYSPORUM 1 (RFO1; also called WALL-ASSOCIATED KINASE-LIKE 22) ^28^ and to a lower extent RECEPTOR-LIKE PROTEIN 44 (RLP44) ^29^ are partially insensitive to 2F-Fuc (Figures 3 A-I and S4 A-I). Our screening also included a mutant for THESEUS 1 (THE1) ^30^ that was previously shown to be involved in cell wall integrity sensing during dark-grown hypocotyl elongation^30^ but did not appear insensitive to 2F-Fuc (Figures 3 A-I and S4 A-I). This finding suggests that both *RFO1* and *RLP44* may be specifically involved in sensing cell wall integrity defect induced by RG-II dimerization deficiency and trigger downstream responses leading to the observed phenotype.

**Figure 3.**
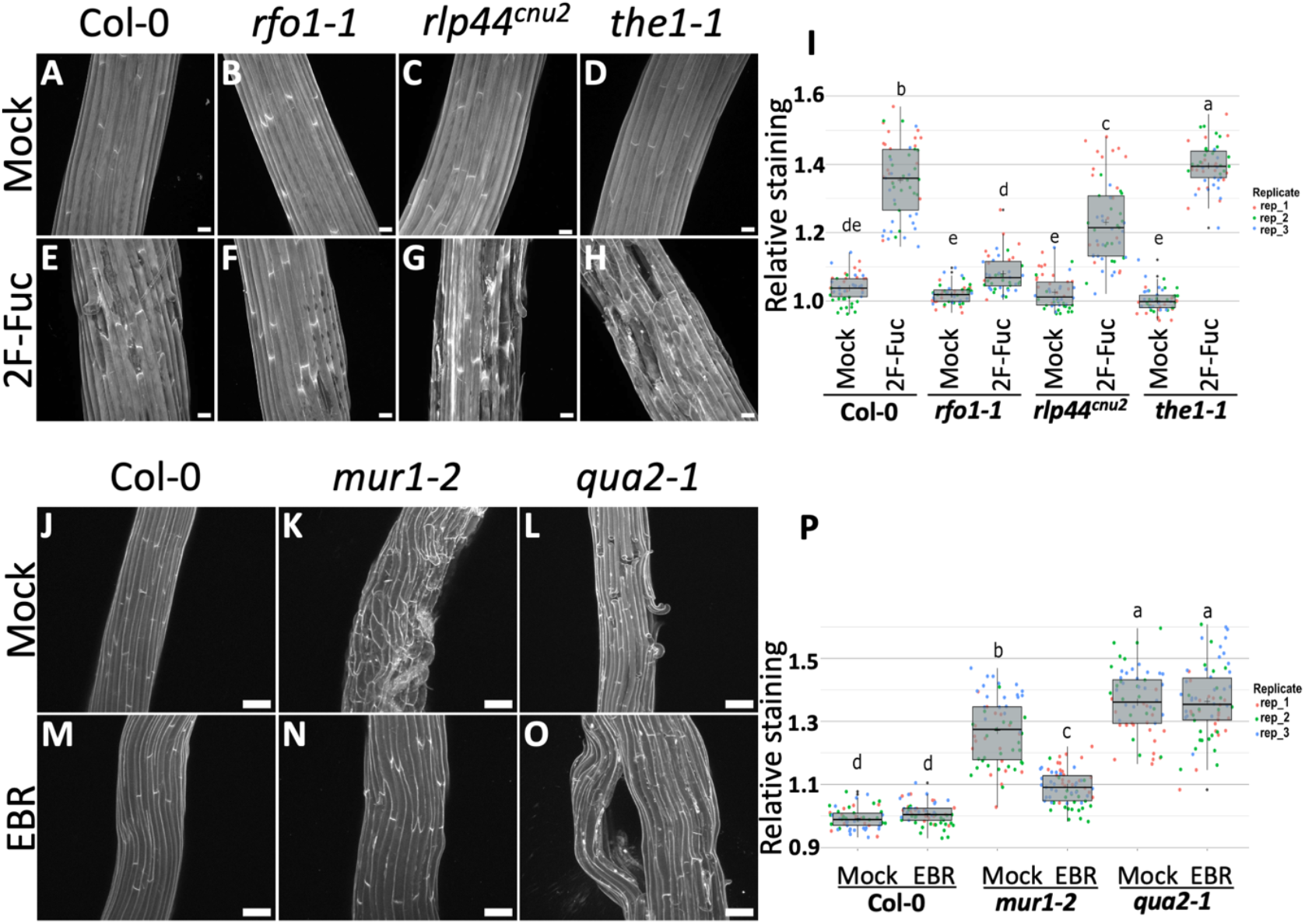
Cell wall sensor mutants and brassinosteroid supplementation rescue RGII dimerization deficiency-induced adhesion defects. (A-O and J-L) Representative max intensity Z-projections of confocal stacks from 4-days-old dark-grown hypocotyls stained with propidium iodide for Col-0 (A, E, J and M), *rfo1-1* (B and F), *rlp44*^*cnu2*^ (C and G), *the1-1* (D and H), *mur1-2* (K and N) and *qua2-1* (L and O), under mock (A-D and J-L), 15µM 2F-Fuc (E-H) and 100nM EBR (M-O) treatment. (I and P) Quantification of the relative ruthenium red staining intensity (see representative images and length quantification in figure S4). Boxplots summarize three biological replicates, as represented by dots of different colors, and letters describe the statistically significant differences between populations determined by one-way ANOVA followed by Tukey’s HSD test (P < 0.05). [Scale bars, 30µm in images (A-H), and 100µm in images (J-O).] See related content in figure S4, S5 and S6.

Both RFO1 and RLP44 were shown to influence brassinosteroid (BR) signalling in response to cell wall integrity defects^28,29^. Thus, we next investigated whether BR also plays a role in our RG-II dimerization deficiency-related adhesion phenotype. We tested the effect of exogenously applied epibrassinolide (EBR; an active form of brassinosteroid) and observed a clear restoration of the adhesion phenotype of *mur1-2* (Figures 3 J-P and S4 J-P). We also performed a q-PCR analysis of BR responsive genes^31,32^ comparing Col-0 and *mur1-2* and found that the BR pathway shows signs of down-regulation in the *mur1* background (Figure S5). Altogether, our results suggest that RG-II dimerization deficiency may be perceived by the cell wall integrity sensors *RFO1* and *RLP44* which may subsequently alter brassinosteroids signalling. Brassinosteroids are known to regulate the expression of genes involved in cell wall synthesis and remodelling^33^. In turn, we can hypothesise that the misregulation of the BR pathway in *mur1* could lead to a weakening of the middle lamella and thus be partially responsible for the observed adhesion defect.

Interestingly, when we performed EBR treatment on *qua2-1* we did not observe a suppression the adhesion defect phenotype (Figures 3 J-P and S4 J-P), further confirming that *QUA2* and *MUR1* mutations affect adhesion through separate pathways. However, we observed a drastic change in the cell separation pattern in *qua2-1* when treated with EBR, where cells separate longitudinally rather than transversely along the hypocotyl (Fig 3 L and O). Furthermore, detached cell files including both epidermal and cortex cells show longitudinal buckling (Fig 3 O) suggesting that the stress pattern leading to cell separation is somehow reversed. Indeed, it has been suggested that transverse cell separation in *qua* mutants was the result of longitudinal tissue tensile stress in the epidermis of highly elongating dark-grown hypocotyls due to the differential and anisotropic growth of inner tissues relative to the growth restricting epidermis^34,35^. Interestingly, BR regulates growth specifically in the epidermal cell layer^36^ and brassinosteroid deficiency has been shown to increase mechanical constrain in the epidermis^37^. Furthermore, mutation of *DWARF4* (BR biosynthesis gene) in the *qua2-1* background was shown to enhance the adhesion defect phenotype by further restricting epidermal growth relative to inner tissues^37^. Conversely, BR is proposed reduce epidermal constrain through cell wall remodelling and thus plays a central role in coordinating growth between cell layers^38^. Thus, while the BR pathway deficiency in *mur1* may limit epidermal growth and increases longitudinal tissue tension, our results on *qua2* also suggest that EBR treatment likely does the opposite by excessively releasing epidermal growth restriction. Cortex cells may drive extensive elongation, unrestricted by the epidermis. In the absence of proper adhesion (in *qua2-1)*, cells become mechanically uncoupled from the rest of the tissues^39^. This uncoupled epidermal and cortex growth may lead to local longitudinal buckling and longitudinal cell detachment of the epidermal and cortex layers. Conversely, the rescue of the *mur1* phenotype by EBR treatment could also be linked to a release of epidermal tension. To test if decreased epidermal tension can be sufficient to rescue *mur1* phenotype, we grew seedlings on medium containing 2.5% agar, (as previously proposed to decrease epidermal tensions)^27,35^, and observed a clear rescue of *mur1* phenotype, similar to the rescue that is observed for *qua2-1* (Figure S6). This suggest that *mur1* adhesion defects can also be linked to improper growth coordination between the epidermis and inner tissues.

To understand the effect of RG-II dimerization deficiency on adhesion and its rescue by EBR, we next characterize the chemistry and mechanics of the cell wall. We first used extensometer-based tensile tests to measure hypocotyl elasticity (tensile modulus) and breaking strength (ultimate tensile strength). We observed that *mur1* hypocotyls have a decreased stiffness and strength compared to Col-0 (Figure 4 A and B), confirming previous observations^40,41^. EBR treatment further decreased stiffness and strength for both Col-0 and *mur1-2* (Figure 4 A and B) further supporting the global effect of BR on cell wall loosening that reduces epidermal tension. On the other hand, the cell wall composition analysis revealed a deficiency in homogalacturonan in *mur1-2* compared to Col-0, which is restored by treatment with EBR (Figure 4 C and S7). These results suggest that *mur1* middle lamella may be weaker than wildtype (lower HG level). In turn, the complementation of *mur1* with EBR could rescue normal adhesion by a combined restoration of normal cell wall composition (restored HG level) and reduced epidermal tension (further suggested here by the decreased tissue stiffness).

**Fig. 4.**
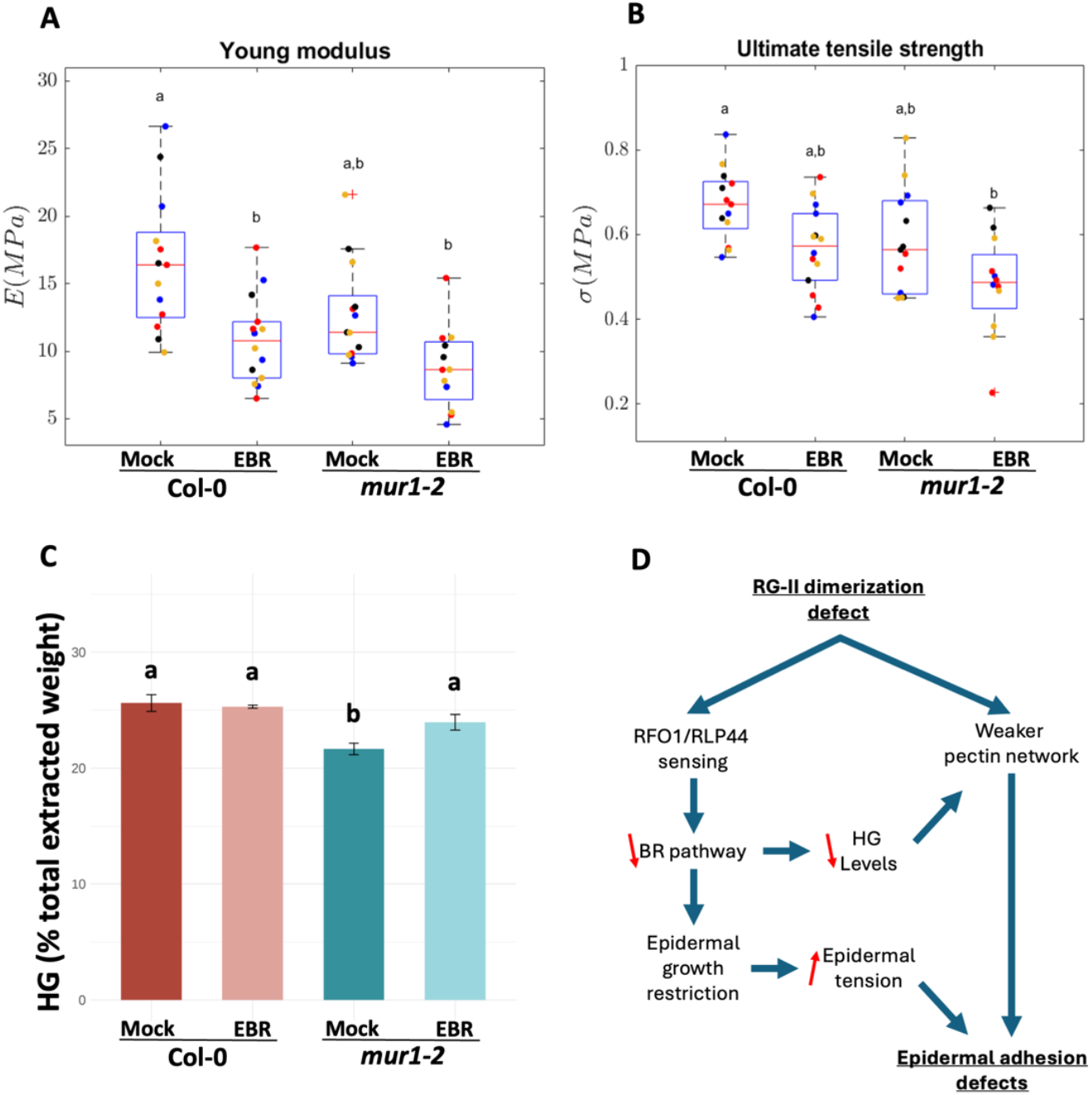
*mur1* mutation and rescue by EBR impact tissue mechanics and homogalacturonan status of the cell wall. (A) Quantification of 4-days-old dark-grown hypocotyl’s Young’s modulus *E*, (B) ultimate tensile strength σand (C) homogalacturonan content (quantity as a percentage of total extracted monosaccharide weight, see related content in figure S7), for Col-0 and *mur1-2* under mock or EBR treatment. Boxplots in (A and B) summarize three biological replicates, as represented by dots of different colors and data in (C) are represented as mean ± SEM. Letters in panels A-C describe the statistically significant differences between different populations determined in (A and B) by one-way ANOVA followed by Tukey-Kramer test (P<0.05) and in (C) by one-way ANOVA followed by Tukey’s HSD test (P < 0.05). See related content in figure S7. (D) Working model for the mechanism of RG-II dimerization defect impact on cell adhesion and integrity: RGII dimerization defects may have both a direct mechanical effect related to the reduction of RGII crosslink, but also affect the structure of the pectin network. RFO1 and RLP44 may sense this pectin integrity defect and tune-down the brassinosteroid (BR) signaling pathway. Miss regulation of the BR pathway may have a dual effect. On one hand impacting homogalacturonan levels and thus likely the middle lamella. On the other hand, by modulating cell layer growth coordination through epidermal growth restriction in turn increasing epidermal tension. The combination of weakened middle lamella and increased epidermal tension would ultimately lead to the observed adhesion defects.

RG-II is a complex component of cell wall whose role in growth and development is not well understood in comparison with cellulose, hemicellulose or homogalacturonan. Our results now identify a role of RG-II dimerization in cell adhesion. Importantly, by studying the effect of reduced RG-II dimerization in plants, we uncover a new pathway regulating the maintenance of cell-cell adhesion. Moreover, we identify the key contribution of cell wall integrity sensing and hormonal regulatory players for the maintenance of cell adhesion in plants mediated by RG-II dimerization. Previous work had hypothesised that such mechanisms should be in place to allow a coordinated regulation of fast extensive growth with the maintenance of adhesion^1,3,35^, but the components and mechanisms had remained mostly unknown. Here, we address this significant gap in our knowledge by identifying the wall associated receptor-like kinase RFO1, the receptor-like protein RLP44 and the brassinosteroid signalling pathway that play a crucial role in coordinating cell wall integrity, cell adhesion and growth coordination. Taken together, our experiments point to a model where RG-II dimerization deficiency leads to cell wall perturbations that are first sensed by the cell wall integrity sensors RFO1 and RLP44. This response modulates the brassinosteroid pathway, leading on one hand to lower HG levels in the middle lamella, and to increased epidermal tension. The combination of the cell-level weakening of the middle lamella, and tissue-level increase in epidermal tension presumably leads to the loss of cell adhesion phenotype observed in response to RG-II dimerization deficiency. Yet, decrease in BR signalling alone cannot fully explain the phenotype observed since down-regulation of the BR pathway alone does not cause adhesion defects on its own. In turn, RG-II crosslinks are also likely to play a direct structural role in the cell wall for adhesion strength. Interestingly this new pathway appears to be largely independent from the previously characterized pathway involving the *quasimodo* and *esmeralda1* mutants. It suggests that both HG and RG-II crosslinks play important roles in the cell wall either directly, mediating adhesive crosslinks, and indirectly as a wall integrity signal regulating downstream response. Here we show that RG-II dimerization defect impacts the BR pathway downstream. Because of the key role of BR in growth control, we propose that normal levels of RG-II dimerization are important to ensure proper feedback signalling contributing to the coordination between fast growth and cell adhesion maintenance in plants. Future work integrating a deeper investigation of the wall integrity defect perception by RFO1 and RLP44 in response to RG-II dimerization deficiency, as well as the downstream perturbation of the BR pathway will surely open the door for groundbreaking discoveries regarding cell wall integrity sensing and cell wall homeostasis mechanisms. Our work provides novel lines of evidence for the complex regulation of cell adhesion in plants, showing that this fundamental feature of multicellular organism indeed necessitates a tight regulation by sensing mechanism and coordination with growth regulators to ensure multicellular tissue integrity.

## Resource availability

### Lead contact

Requests for further information and resources should be directed to and will be fulfilled by the lead contact, Stephane Verger

### Materials availability

All material generated in this study will be made available upon request to the lead contact.

### Data and code availability

#### Data

- All the confocal microscopy data used to support this study (including those not displayed in figures) has been deposited at Zenodo and is publicly available as of the date of publication (10.5281/zenodo.14170979).
- All the light microscopy data for ruthenium red staining experiments along with the processed data (including segmented masks, corrected masks, raw quantifications, processed quantifications) has been deposited at Zenodo and is publicly available as of the date of publication (10.5281/zenodo.14171190).
- All the light microscopy and measurements data for the tensile test experiment has been deposited at Zenodo and is publicly available as of the date of publication (10.5281/zenodo.14171222).
- Raw data for the Q-PCR analysis and cell wall quantification reported in this paper will be shared by the lead contact upon request.

#### Code

- The code for the ruthenium red quantification workflow is publicly available on github (github.com/VergerLab/RRQuant) and the specific version used for this study has been archived at Zenodo and is publicly available as of the date of publication at (10.5281/zenodo.14173185).
- The code for the extensometer control and data analysis has been deposited at Zenodo and is publicly available as of the date of publication (10.5281/zenodo.14171222).

#### Additional information

- Any additional information required to reanalyze the data reported in this paper is available from the lead contact upon request.

## Supporting information

Data S1- Related to Figures 1-4 and S2-S7

## Acknowledgment

We thank Olivier Hamant, Grégory Mouille and Stéphanie Robert for helpful comments on the manuscript. We thank Clara Sánchez-Rodríguez, Sebastian Wolf, Paul Dupree, Richard Stresser, Tai-ping Sun and Julien Sechet for providing the seeds of *rfo1-1, rlp44*^*cnu2*^, *fut4* and *fut4/6, FucTA/FucTB, spy-8* and *spy-12* and hp*GGLT1* lines. We thank the facilities and technical assistance of the Umeå Plant Science Centre (UPSC) microscopy facility, biopolymer analytical platform and the plant growth facility. We thank Sarah Robinson and Mateusz Majda for initial help setting up our micro extensometer, Elsa Demes for initial help setting up the ruthenium red staining and high-throughput imaging method and Kaël Lissarrague for other technical help.

This work was supported by grants from the Swedish research council (VR, 2020– 03974), Novo Nordisk foundation (NNF21OC0067282), Åforsk foundation (20-502), Carl Kempe foundation (JCK-1912.2) and Carl Tryggers foundation (CTS19: 398) to S.V. P.J.K. and B.A. were funded by postdoc fellowship from Beijing Advanced center for molecular design. R.P.B. was funded by grants from VR and HFSP. R.S.S. was supported by a Biotechnological and Biological Sciences Research Council (BBSRC) Institute Strategic Program Grant (BB/X01102X/1) to the John Innes Centre. This work was also supported by Umeå Plant Science Centre with grants from the Knut and Alice Wallenberg Foundation (KAW 2016.0341 and KAW 2016.0352), the Swedish Governmental Agency for Innovation Systems (VINNOVA 2016–00504) and Bio4Energy, a Strategic Research Environment supported through the Swedish Government’s Strategic Research Area initiative.

## Author contributions

Conceptualization, S.V., R.P.B and A.I.B; Methodology, A.I.B., L.L., A.A., L.B., O.E., A.H. and S.V.; Software, A.A., L.B., A.H., R.S.S. and S.V.; Formal Analysis, A.I.B., L.L., A.A. and L.B.; Investigation, A.I.B., L.L. and A.A.; Resources, B.A. and P.J.K.; Writing – Original Draft, A.I.B. and S.V.; Writing – Review & Editing, A.I.B., L.L., A.A., L.B., O.E., A.H., S.V., R.S.S. and R.P.B.; Visualization, A.I.B., L.L., A.A., L.B. and S.V.; Supervision, S.V. and R.P.B.; Funding Acquisition, S.V., R.P.B. and R.S.S.

## Supplemental information

Document S1. Figures S1-S7

Data S1. Excel file containing the source data for the experiments in this study too large to fit in a PDF, related to Figures 1-4 and S2–S7.

## Methods

### Plant growth conditions

Plants were grown on 1/2 MS (Murashige-Skoog) medium (Duchefa) which also contained 0.5 % (w/v) sucrose, 0.8% plant agar (Duchefa). The pH of the media was set to 5.7 by NaOH which was buffered by using 0.05% (w/v) 2-(N-morpholino)ethanesulfonic acid (MES) (Sigma-Aldrich). Sterilized seeds were then put on media and stratified at 4°C for 2 days in darkness and then light induced for 6 hours in white light at 21°C. After the light induction the petri plates containing the seeds were wrapped in few layers of aluminum foil and left vertically to germinate in the dark at 21°C for 4 days. In case of high agar experiments, 2.5% plant agar was used in the media and 0.8% plant agar was used as control media, with other constituents in media kept similar. In experiments with liquid media the constituents were same as above but did not contain any agar.

### Plant materials

*Arabidopsis thaliana* ecotype Columbia (Col-0) was used as wildtype control in all experiments with the exception of one experiment where Landsberg erecta (Ler) was used for a comparison with a mutant in the Ler background. The mutant lines *mur1-1* and *mur1-2* ^18,42^, *qua2-1* ^9^, *esmd1-1* and *qua2-1 esmd1-1* ^3^, *mur2-1* ^19,43^, *fut4* and *fut4/6* ^20^, *FucTA/FucTB* ^21^, *spy8* and *spy12* ^22,23^, *rfo1-1* ^28^, *rlp44*^*cnu2* 29^, *the1-1* ^30^, hp*GGLT1* (Line #1 and Line #3) RNAi transgenic lines with a hairpin (hp) RNA construct ^26^ have been described earlier. We also screened the mutant lines *fer-4* ^44^, *herk2-1* and *herk1-1* ^45^, *cvy1-1* ^46^ and *wakΔ5* ^13^. The double mutants *mur1-2 esmd1-1* and *mur1-2 qua2-1* have been generated in this study. Primers used in the genotyping of the mutants were those described in the corresponding references, except for the *rfo1-1* left primer for which we used TCACCGTAAACCCATTAG.

### Chemical treatments

For fucosylation inhibition, 2-Deoxy-2-fluoro-L-fucose (2F-Fuc) (MedChemExpress) dissolved in DMSO by ultrasonication and then filter sterilized was used. Sterilized seeds were grown on ½ MS liquid media containing 15µM 2F-Fuc. For boron treatment, boric acid (Sigma) prepared by dissolving in MilliQ water and then filter sterilized was used. Sterilized seeds were grown on ½ MS solid media containing either 500µM or 750µM of boric acid. For brassinosteroid treatments, epibrassinolide (EBR) (Sigma-Aldrich) dissolved in DMSO and filter sterilized was used. Sterilized seeds were grown on ½ MS solid media containing 100nM EBR. For all treatments, seedlings were grown for 4 days in the dark as described in the “plant growth condition” section and DMSO was used as mock control in media except for boric acid treatment.

### Propidium iodide staining and confocal microscopy

Dark-grown seedlings were first immersed for 5 minutes in 0.1mg/ml propidium iodide (Sigma-Aldrich) prepared in milliQ water and then they were washed with water before imaging. Seedlings were then placed between glass slides and coverslip with spacers in between them to prevent sample crushing. Images of hypocotyls were taken with a Zeiss LSM 880 inverted laser scanning confocal microscope using the Zen blue software for acquisition. Samples were imaged with either 10X (air NA 0.3) or 32X (Water, NA 0.85) objectives. Z-stacks were acquired without averaging and with a 0.5 µm Z step. The images have a size of 1024 × 1024px for a pixel size of 0.83 or 0.42 µm. Excitation of PI was performed by using a 552 nm solid-state laser and the fluorescence was detected at 600–650 nm. A minimum of 7 seedlings were imaged for each genotype and conditions and one representative image was included in the figures.

### Ruthenium Red staining and dark field microscopy

Four-days-old dark-grown seedlings were immersed for 1 minute in 0.05% (w/v) ruthenium red (Sigma-Aldrich) solution prepared in milliQ water and the seedlings were immediately washed in water. Washed seedlings were then transferred on square petri plates containing 50 ml of solidified 1% agarose along with two or three non-stained seedlings per genotype and conditions. A Leica M205CFA microscope with 1X/0.02NA objective was used to acquire large tile RGB darkfield images of the plates. Large, stitched tile images were generally made of 90 to 110 individual 1920×1080px images with a pixel size of 6.29 µm that were acquired with 10% overlap and merged with the Leica Application Suite (LAS X) software. Experiments were done in three independent biological replicates with 15 or more seedlings for each genotype or treatment per replicate.

### Quantification of ruthenium red staining and hypocotyl length

Ruthenium red staining intensity was quantified in hypocotyls with a workflow involving the automated deep learning-based hypocotyl segmentation followed by quantification of relative staining intensity, hypocotyl morphology and statistical analysis (github.com/VergerLab/RRQuant). Briefly, we used root painter^47^ to train a U-Net CNN deep learning model for high-throughput and accurate hypocotyl segmentation from our darkfield images of stained and non-stained samples. If needed segmentation masks were further manually corrected and saved using a Fiji ^48^ macro. We then developed another Fiji macro for the quantification of staining intensity and hypocotyl morphology. Ruthenium red staining intensity was calculated by converting the RGB image into an HSB (Hue, Saturation, Brightness) stack, inverting the “brightness” image values, and creating a new “intensity” image by adding the “saturation” and inverted “brightness” pixel values. Staining intensity was quantified as the average pixel value in the “intensity” image within the region of the segmented mask that was eroded by 10px to limit edge coloration effects, for each hypocotyl individually. Hypocotyl length was quantified using the geodesic diameter measurement values of the MorpholibJ ImageJ plugin ^49^ on the non-eroded masks. We then developed a script for data processing, plot generation and statistical analysis in R (https://www.R-project.org). Relative staining intensity was calculated as the ratio of individual stained samples over the average value measured on non-stained reference samples of the corresponding genotype and condition, treating each replicates independently. Relative staining intensity and hypocotyl length values were used to generate plots in R studio. Statistically significant differences among the samples were determined by one-way ANOVA followed by the Tukey’s HSD (post hoc) test (P < 0.05) or by two-tailed Student’s t test wherever applicable.

### RNA isolation and quantitative Real-Time PCR analysis

RNA isolation was performed on the 4-days-old dark-grown hypocotyls from the control and mutant seedlings. Sampling of the seedings was performed under green light and then were immediately frozen in the liquid nitrogen. The total RNA of each sample was extracted using the RNeasy Plant Mini Kit (QIAGEN). Following extraction, total RNA was treated with RNase-free DNase (Thermo Fisher Scientific). cDNA was synthesized using iScript™ cDNA Synthesis Kit (Bio-Rad) and was used in quantitative real-time PCR (qRT–PCR). qRT-PCR was carried out using iTaq Universal SYBR Green Supermix (Bio-Rad) according to the manufacturer’s instruction. Primers used for the qRT-PCR studies are those described in ^31,32^. The data was analyzed using the method described in ^50,51^. UBQ10 and ACT2 were used as two reference genes. Plots and the statistical analysis were done using RStudio. Statistically significant difference between the samples was determined by a two-tailed Student’s t test (P<0.05).

### Young modulus and ultimate tensile strength measurements

Our micro-extensometer setup is similar to the ones used previously^52–54^ that use SmarAct positioners (MCS2, SLC-1720) to stretch the sample and Futek load cells (Futek LSB200) to measure force. To image samples during the experiments, we used an upright brightfield imaging setup that consists of a color CMOS camera (IDS; U3-3080SE-C-HQ R), a 12X zoom Lens (Thorlabs; MVL12×3Z) and white light transmission illumination. We developed software in the MorphRobotX (MRX) framework of MorphoDynamX (morphographx.org/morphorobotx/) to synchronize image capture from our setup with extensometer steps. This allowed us to use landmarks on the images to track the deformation during stretching, and avoid artifacts due to slippage of the sample at the anchoring sites. Four-day-old dark-grown hypocotyls were mounted on the extensometer arms. This was done using two pieces of tough-tag (Merk; Z359106) placed parallel to each other with approximately 2mm of spacing in between. A small piece of silicone transfer adhesive (Adhesive Applications; PolySil™ S5005DC; 1×4mm) was placed at the edge of each tough tags. Individual hypocotyls (one per stretching measurement) were placed in between the two silicon adhesives. Another piece of the silicon adhesive was placed on top of the hypocotyl in a way that the samples is sandwiched between the silicon adhesive tapes. The tough-tags were then transferred to the arms of the micro extensometer before the arms of the extensometer were lowered into a petri dish filled with Milli-Q water. Extensometer experiments were automatically performed with MRX (processes/Tools/MorphoRobotX/Experiment/Extensometer). The stretching was performed at a constant speed of 7.7µm/s with the nanopositionners of the extensometer moving 1µm in each measurement step. Images acquired with the camera were used to estimate the sample cross section area and the strain during stretching. For cross section area, we used Fiji to draw three lines across the width of the hypocotyl at three different positions on the image at the first time point (before stretching). The average width value (w) was used to calculate the cross-section area (A) by approximating the hypocotyl as cylinder (π*(w/2)^2^). For strain quantification, a MATLAB code was created to extract and track two points on the hypocotyl in the video recorded during stretching. The distance between these two points at the first time point (L0) and throughout the experiment (L) was quantified in pixels and used to calculate the strain ((L-L0)/L0) at each time point. The value from the load cell was used to calculate the stress (F/A) at each time point as the force (F) divided by the initial circular cross-section area (A) of the sample. The stress and strain values were used to plot stress-strain curves for each stretched samples and calculate the Young modulus and ultimate tensile strength. The Young modulus of the hypocotyl was extracted by fitting a line between two points in the linear part of the stress-strain curve and using the slope as the elasticity value. The ultimate tensile strength was extracted from the curves as the maximum stress imposed to the sample. We ran four independent biological replicates, measuring 4 to 5 samples per genotype and treatment in each replicate, thus generating 16 to 20 measurements per genotype and treatment. Several measurements were excluded due to visible issue with sample slippage, abnormalities in the trend of the stress-strain curve or absence of identifiable linear regime, reducing to 12-13 samples per genotypes and treatments being included in the final analysis and plots. Data analysis and graphs generation was done in MATLAB. Statistically significant difference between the samples was determined by one-way ANOVA followed by Tukey-Kramer test (P<0.05).

### Cell wall analysis

Whole four-days-old dark-grown seedlings were freeze-dried and subsequently ground using a ball-mill. To obtain the alcohol insoluble residue (AIR), 10 mg of grounded material per sample was first incubated for 30 min at RT with 80% EtOH followed by chloroform: methanol (1:1) incubation for another 30 min. The material was dried and 1000 units of α-amylase in 0.1M potassium phosphate buffer was added to remove the starch. After overnight incubation, the samples were first washed with Milli-Q water and then with acetone and left to dry in the vacuum desiccator overnight. 0.5 mg DW of previously treated samples was taken for monosaccharide analysis by Trimethylsilyl derivatization (TMS) and were analysed as described previously^55^ with GC/MS (7890A/5975C; Agilent Technologies). To determine the degree of methyl esterification, 1.5 mg DW was taken from the AIR treated samples. The material was first saponified by incubating it with 1M NaOH for 1h at RT after which the reaction was neutralized with 1M HCl. Around 180 µL of supernatant was taken for further methanol detection. Reactions were prepared in ELISA-plates where 50 µL of alcohol oxidase in 0.1M sodium phosphate buffer was added to each sample and was incubated for 15 min at RT. After that, 100 µL of developer containing acetylacetone, acetic acid and ammonium acetate in water was added in each sample, covered and incubated at 68°C for 10 min. Finally, the absorption was read at 412 nm. The analysis was done as previously described^56^. The homogalacturonan content was approximated by subtracting the quantified rhamnose from galacturonic acid values. Data analysis and graphs generation was done in RStudio. Statistically significant difference between the samples was determined by one-way ANOVA followed by Tukey HSD test (P<0.05), Welch’s ANOVA followed by Games-Howell test (P<0.05) and Kruskal-Wallis with Dunn’s test for pairwise comparisons (P<0.05) where applicable.

## References

1. Baba, A.I., and Verger, S. (2024). Cell adhesion maintenance and controlled separation in plants. Front. Plant Physiol. 2, 1369575. 10.3389/fphgy.2024.1369575.

2. Anderson, C.T. (2019). Pectic Polysaccharides in Plants: Structure, Biosynthesis, Functions, and Applications. In Extracellular Sugar-Based Biopolymers Matrices Biologically-Inspired Systems., E. Cohen and H. Merzendorfer, eds. (Springer International Publishing), pp. 487–514. 10.1007/978-3-030-12919-4_12.

3. Verger, S., Chabout, S., Gineau, E., and Mouille, G. (2016). Cell adhesion in plants is under the control of putative O-fucosyltransferases. Development, dev. 132308. 10.1242/dev.132308.

4. Knox, J.P. (1992). Cell adhesion, cell separation and plant morphogenesis. The Plant Journal 2, 137–141. 10.1111/j.1365-313X.1992.00137.x.

5. Jarvis, M.C., Briggs, S.P.H., and Knox, J.P. (2003). Intercellular adhesion and cell separation in plants. Plant Cell & Environment 26, 977–989. 10.1046/j.1365-3040.2003.01034.x.

6. Daher, F.B., and Braybrook, S.A. (2015). How to let go: pectin and plant cell adhesion. Front. Plant Sci. 6. 10.3389/fpls.2015.00523.

7. Atakhani, A., Bogdziewiez, L., and Verger, S. (2022). Characterising the mechanics of cell–cell adhesion in plants. Quant Plant Bio. 3, e2. 10.1017/qpb.2021.16.

8. Bouton, S., Leboeuf, E., Mouille, G., Leydecker, M.-T., Talbotec, J., Granier, F., Lahaye, M., Höfte, H., and Truong, H.-N. (2002). QUASIMODO1 Encodes a Putative Membrane-Bound Glycosyltransferase Required for Normal Pectin Synthesis and Cell Adhesion in Arabidopsis. Plant Cell 14, 2577–2590. 10.1105/tpc.004259.

9. Mouille, G., Ralet, M., Cavelier, C., Eland, C., Effroy, D., Hématy, K., McCartney, L., Truong, H.N., Gaudon, V., Thibault, J., et al. (2007). Homogalacturonan synthesis in Arabidopsis thaliana requires a Golgi-localized protein with a putative methyltransferase domain. The Plant Journal 50, 605–614. 10.1111/j.1365-313X.2007.03086.x.

10. Du, J., Kirui, A., Huang, S., Wang, L., Barnes, W.J., Kiemle, S.N., Zheng, Y., Rui, Y., Ruan, M., Qi, S., et al. (2020). Mutations in the Pectin Methyltransferase QUASIMODO2 Influence Cellulose Biosynthesis and Wall Integrity in Arabidopsis. The Plant Cell 32, 3576–3597. 10.1105/tpc.20.00252.

11. Barnes, W.J., Zelinsky, E., and Anderson, C.T. (2022). Polygalacturonase activity promotes aberrant cell separation in the quasimodo2 mutant of Arabidopsis thaliana. The Cell Surface 8, 100069. 10.1016/j.tcsw.2021.100069.

12. Grandjean, C., Voxeur, A., Chabout, S., Jobert, F., Gutierrez, L., Pelloux, J., Mouille, G., and Bouton, S. (2024). Fine-tuning and remodeling of pectins play a key role in the maintenance of cell adhesion. Front. Plant Physiol. 2, 1441158. 10.3389/fphgy.2024.1441158.

13. Kohorn, B.D., Greed, B.E., Mouille, G., Verger, S., and Kohorn, S.L. (2021). Effects of Arabidopsis wall associated kinase mutations on ESMERALDA1 and elicitor induced ROS. PLoS ONE 16, e0251922. 10.1371/journal.pone.0251922.

14. Iwai, H., Masaoka, N., Ishii, T., and Satoh, S. (2002). A pectin glucuronyltransferase gene is essential for intercellular attachment in the plant meristem. Proc. Natl. Acad. Sci. U.S.A. 99, 16319–16324. 10.1073/pnas.252530499.

15. Voxeur, A., Soubigou-Taconnat, L., Legée, F., Sakai, K., Antelme, S., Durand-Tardif, M., Lapierre, C., and Sibout, R. (2017). Altered lignification in mur1-1 a mutant deficient in GDP-L-fucose synthesis with reduced RG-II cross linking. PLoS ONE 12, e0184820. 10.1371/journal.pone.0184820.

16. Waszczak, C., Yarmolinsky, D., Leal Gavarrón, M., Vahisalu, T., Sierla, M., Zamora, O., Carter, R., Puukko, T., Sipari, N., Lamminmäki, A., et al. (2024). Synthesis and import of GDP-L -fucose into the Golgi affect plant–water relations. New Phytologist 241, 747–763. 10.1111/nph.19378.

17. O’Neill, M.A., Eberhard, S., Albersheim, P., and Darvill, A.G. (2001). Requirement of Borate Cross-Linking of Cell Wall Rhamnogalacturonan II for Arabidopsis Growth. Science 294, 846–849. 10.1126/science.1062319.

18. Bonin, C.P., Potter, I., Vanzin, G.F., and Reiter, W.-D. (1997). The MUR1 gene of Arabidopsis thaliana encodes an isoform of GDP- D -mannose-4,6-dehydratase, catalyzing the first step in the de novo synthesis of GDP- L -fucose. Proc. Natl. Acad. Sci. U.S.A. 94, 2085–2090. 10.1073/pnas.94.5.2085.

19. Reiter, W., Chapple, C., and Somerville, C.R. (1997). Mutants of Arabidopsis thaliana with altered cell wall polysaccharide composition. The Plant Journal 12, 335–345. 10.1046/j.1365-313X.1997.12020335.x.

20. Tryfona, T., Theys, T.E., Wagner, T., Stott, K., Keegstra, K., and Dupree, P. (2014). Characterisation of FUT4 and FUT6 α-(1→2)-Fucosyltransferases Reveals that Absence of Root Arabinogalactan Fucosylation Increases Arabidopsis Root Growth Salt Sensitivity. PLoS ONE 9, e93291. 10.1371/journal.pone.0093291.

21. Strasser, R., Altmann, F., Mach, L., Glössl, J., and Steinkellner, H. (2004). Generation of Arabidopsis thaliana plants with complex N -glycans lacking β1,2-linked xylose and core α1,3-linked fucose. FEBS Letters 561, 132–136. 10.1016/S0014-5793(04)00150-4.

22. Silverstone, A.L., Tseng, T.-S., Swain, S.M., Dill, A., Jeong, S.Y., Olszewski, N.E., and Sun, T. (2007). Functional Analysis of SPINDLY in Gibberellin Signaling in Arabidopsis. Plant Physiology 143, 987–1000. 10.1104/pp.106.091025.

23. Zentella, R., Sui, N., Barnhill, B., Hsieh, W.-P., Hu, J., Shabanowitz, J., Boyce, M., Olszewski, N.E., Zhou, P., Hunt, D.F., et al. (2017). The Arabidopsis O-fucosyltransferase SPINDLY activates nuclear growth repressor DELLA. Nat Chem Biol 13, 479–485. 10.1038/nchembio.2320.

24. Dumont, M., Lehner, A., Bardor, M., Burel, C., Vauzeilles, B., Lerouxel, O., Anderson, C.T., Mollet, J., and Lerouge, P. (2015). Inhibition of fucosylation of cell wall components by 2-fluoro 2-deoxy- L -fucose induces defects in root cell elongation. The Plant Journal 84, 1137–1151. 10.1111/tpj.13071.

25. Villalobos, J.A., Yi, B.R., and Wallace, I.S. (2015). 2-Fluoro-L-Fucose Is a Metabolically Incorporated Inhibitor of Plant Cell Wall Polysaccharide Fucosylation. PLoS ONE 10, e0139091. 10.1371/journal.pone.0139091.

26. Sechet, J., Htwe, S., Urbanowicz, B., Agyeman, A., Feng, W., Ishikawa, T., Colomes, M., Kumar, K.S., Kawai-Yamada, M., Dinneny, J.R., et al. (2018). Suppression of Arabidopsis GGLT 1 affects growth by reducing the L-galactose content and borate cross-linking of rhamnogalacturonan-II. The Plant Journal 96, 1036–1050. 10.1111/tpj.14088.

27. Malivert, A., Erguvan, Ö., Chevallier, A., Dehem, A., Friaud, R., Liu, M., Martin, M., Peyraud, T., Hamant, O., and Verger, S. (2021). FERONIA and microtubules independently contribute to mechanical integrity in the Arabidopsis shoot. PLoS Biol 19, e3001454. 10.1371/journal.pbio.3001454.

28. Huerta, A.I., Sancho-Andrés, G., Montesinos, J.C., Silva-Navas, J., Bassard, S., Pau-Roblot, C., Kesten, C., Schlechter, R., Dora, S., Ayupov, T., et al. (2023). The WAK-like protein RFO1 acts as a sensor of the pectin methylation status in Arabidopsis cell walls to modulate root growth and defense. Molecular Plant 16, 865–881. 10.1016/j.molp.2023.03.015.

29. Wolf, S., Van Der Does, D., Ladwig, F., Sticht, C., Kolbeck, A., Schürholz, A.-K., Augustin, S., Keinath, N., Rausch, T., Greiner, S., et al. (2014). A receptor-like protein mediates the response to pectin modification by activating brassinosteroid signaling. Proc. Natl. Acad. Sci. U.S.A. 111, 15261–15266. 10.1073/pnas.1322979111.

30. Hématy, K., Sado, P.-E., Van Tuinen, A., Rochange, S., Desnos, T., Balzergue, S., Pelletier, S., Renou, J.-P., and Höfte, H. (2007). A Receptor-like Kinase Mediates the Response of Arabidopsis Cells to the Inhibition of Cellulose Synthesis. Current Biology 17, 922–931. 10.1016/j.cub.2007.05.018.

31. Bittner, T., Nadler, S., Schulze, E., and Fischer-Iglesias, C. (2015). Two homolog wheat Glycogen Synthase Kinase 3/SHAGGY - like kinases are involved in brassinosteroid signaling. BMC Plant Biol 15, 247. 10.1186/s12870-015-0617-z.

32. Thussagunpanit, J., Jutamanee, K., Homvisasevongsa, S., Suksamrarn, A., Yamagami, A., Nakano, T., and Asami, T. (2017). Characterization of synthetic ecdysteroid analogues as functional mimics of brassinosteroids in plant growth. The Journal of Steroid Biochemistry and Molecular Biology 172, 1–8. 10.1016/j.jsbmb.2017.05.003.

33. Sun, Y., Fan, X.-Y., Cao, D.-M., Tang, W., He, K., Zhu, J.-Y., He, J.-X., Bai, M.-Y., Zhu, S., Oh, E., et al. (2010). Integration of Brassinosteroid Signal Transduction with the Transcription Network for Plant Growth Regulation in Arabidopsis. Developmental Cell 19, 765–777. 10.1016/j.devcel.2010.10.010.

34. Kutschera, U., and Niklas, K.J. (2007). The epidermal-growth-control theory of stem elongation: An old and a new perspective. Journal of Plant Physiology 164, 1395–1409. 10.1016/j.jplph.2007.08.002.

35. Verger, S., Long, Y., Boudaoud, A., and Hamant, O. (2018). A tension-adhesion feedback loop in plant epidermis. eLife 7, e34460. 10.7554/eLife.34460.

36. Savaldi-Goldstein, S., Peto, C., and Chory, J. (2007). The epidermis both drives and restricts plant shoot growth. Nature 446, 199–202. 10.1038/nature05618.

37. Kelly-Bellow, R., Lee, K., Kennaway, R., Barclay, J.E., Whibley, A., Bushell, C., Spooner, J., Yu, M., Brett, P., Kular, B., et al. (2023). Brassinosteroid coordinates cell layer interactions in plants via cell wall and tissue mechanics. Science 380, 1275–1281. 10.1126/science.adf0752.

38. Aardening, Z., Khandal, H., Erlichman, O.A., and Savaldi-Goldstein, S. (2024). The whole and its parts: cell-specific functions of brassinosteroids. Trends in Plant Science, S1360138524002838. 10.1016/j.tplants.2024.10.015.

39. Verger, S., Liu, M., and Hamant, O. (2019). Mechanical Conflicts in Twisting Growth Revealed by Cell-Cell Adhesion Defects. Front. Plant Sci. 10, 173. 10.3389/fpls.2019.00173.

40. Abasolo, W., Eder, M., Yamauchi, K., Obel, N., Reinecke, A., Neumetzler, L., Dunlop, J.W.C., Mouille, G., Pauly, M., Höfte, H., et al. (2009). Pectin May Hinder the Unfolding of Xyloglucan Chains during Cell Deformation: Implications of the Mechanical Performance of Arabidopsis Hypocotyls with Pectin Alterations. Molecular Plant 2, 990–999. 10.1093/mp/ssp065.

41. Ryden, P., Sugimoto-Shirasu, K., Smith, A.C., Findlay, K., Reiter, W.-D., and McCann, M.C. (2003). Tensile Properties of Arabidopsis Cell Walls Depend on Both a Xyloglucan Cross-Linked Microfibrillar Network and Rhamnogalacturonan II-Borate Complexes. Plant Physiology 132, 1033–1040. 10.1104/pp.103.021873.

42. Reiter, W.-D., Chapple, C.C.S., and Somerville, C.R. (1993). Altered Growth and Cell Walls in a Fucose-Deficient Mutant of Arabidopsis. Science 261, 1032–1035. 10.1126/science.261.5124.1032.

43. Vanzin, G.F., Madson, M., Carpita, N.C., Raikhel, N.V., Keegstra, K., and Reiter, W.-D. (2002). The mur2 mutant of Arabidopsis thaliana lacks fucosylated xyloglucan because of a lesion in fucosyltransferase AtFUT1. Proc. Natl. Acad. Sci. U.S.A. 99, 3340–3345. 10.1073/pnas.052450699.

44. Duan, Q., Kita, D., Li, C., Cheung, A.Y., and Wu, H.-M. (2010). FERONIA receptor-like kinase regulates RHO GTPase signaling of root hair development. Proc. Natl. Acad. Sci. U.S.A. 107, 17821–17826. 10.1073/pnas.1005366107.

45. Guo, H., Li, L., Ye, H., Yu, X., Algreen, A., and Yin, Y. (2009). Three related receptor-like kinases are required for optimal cell elongation in Arabidopsis thaliana. Proc. Natl. Acad. Sci. U.S.A. 106, 7648–7653. 10.1073/pnas.0812346106.

46. Gachomo, E.W., Jno Baptiste, L., Kefela, T., Saidel, W.M., and Kotchoni, S.O. (2014). The Arabidopsis CURVY1 (CVY1) gene encoding a novel receptor-like protein kinase regulates cell morphogenesis, flowering time and seed production. BMC Plant Biol 14, 221. 10.1186/s12870-014-0221-7.

47. Smith, A.G., Han, E., Petersen, J., Olsen, N.A.F., Giese, C., Athmann, M., Dresbøll, D.B., and Thorup-Kristensen, K. (2022). R OOT P AINTER : deep learning segmentation of biological images with corrective annotation. New Phytologist 236, 774–791. 10.1111/nph.18387.

48. Schindelin, J., Arganda-Carreras, I., Frise, E., Kaynig, V., Longair, M., Pietzsch, T., Preibisch, S., Rueden, C., Saalfeld, S., Schmid, B., et al. (2012). Fiji: an open-source platform for biological-image analysis. Nat Methods 9, 676–682. 10.1038/nmeth.2019.

49. Legland, D., Arganda-Carreras, I., and Andrey, P. (2016). MorphoLibJ: integrated library and plugins for mathematical morphology with ImageJ. Bioinformatics 32, 3532–3534. 10.1093/bioinformatics/btw413.

50. Vandesompele, J., De Preter, K., Pattyn, F., Poppe, B., Van Roy, N., De Paepe, A., and Speleman, F. (2002). Accurate normalization of real-time quantitative RT-PCR data by geometric averaging of multiple internal control genes. Genome Biol 3, research0034.1. 10.1186/gb-2002-3-7-research0034.

51. Hellemans, J., Mortier, G., De Paepe, A., Speleman, F., and Vandesompele, J. (2007). qBase relative quantification framework and software for management and automated analysis of real-time quantitative PCR data. Genome Biol 8, R19. 10.1186/gb-2007-8-2-r19.

52. Hofhuis, H., Moulton, D., Lessinnes, T., Routier-Kierzkowska, A.-L., Bomphrey, R.J., Mosca, G., Reinhardt, H., Sarchet, P., Gan, X., Tsiantis, M., et al. (2016). Morphomechanical Innovation Drives Explosive Seed Dispersal. Cell 166, 222–233. 10.1016/j.cell.2016.05.002.

53. Robinson, S., Huflejt, M., Barbier De Reuille, P., Braybrook, S.A., Schorderet, M., Reinhardt, D., and Kuhlemeier, C. (2017). An Automated Confocal Micro-Extensometer Enables in Vivo Quantification of Mechanical Properties with Cellular Resolution. Plant Cell 29, 2959–2973. 10.1105/tpc.17.00753.

54. Majda, M., Trozzi, N., Mosca, G., and Smith, R.S. (2022). How Cell Geometry and Cellular Patterning Influence Tissue Stiffness. IJMS 23, 5651. 10.3390/ijms23105651.

55. Sweeley, C.C., Bentley, Ronald., Makita, M., and Wells, W.W. (1963). Gas-Liquid Chromatography of Trimethylsilyl Derivatives of Sugars and Related Substances. J. Am. Chem. Soc. 85, 2497–2507. 10.1021/ja00899a032.

56. Klavons, J.A., and Bennett, R.D. (1986). Determination of methanol using alcohol oxidase and its application to methyl ester content of pectins. J. Agric. Food Chem. 34, 597–599. 10.1021/jf00070a004.

