## Supplementary material for "Rhamnogalacturonan-II dimerization deficiency impairs the coordination between growth and adhesion maintenance in plants": Document S1 - Figures S1-S7

### Figure S1

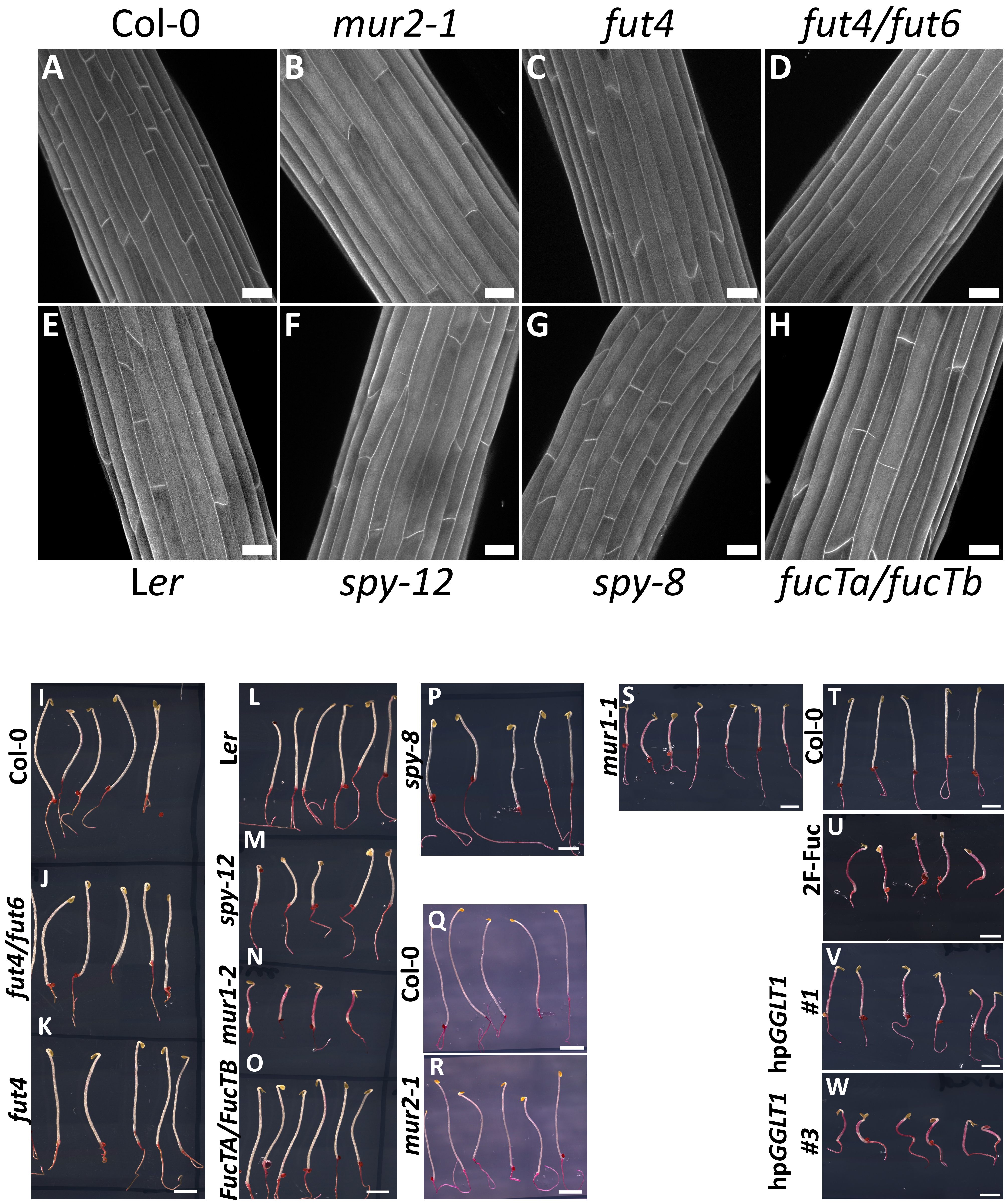


**Fig S1. RG-II dimerization is crucial for cellular and supracellular integrity. Related to figure 1.** (A-H) Representative max intensity Z-projections of the confocal stacks from 4-days-old dark-grown hypocotyl stained with propidium iodide from Col-0 (A) , *mur2-1* (B)*, fut4* (C)*, fut4/fut6* (D)*,* L*er* (E), *spy12* (F)*, spy8* (G) *and FucTA/FucTB* (H). (I-W) Representative images of ruthenium red stained 4-days-old dark-grown hypocotyls for Col-0 (I, Q and T), *fut4/fut6* (J)*, fut4* (K)*,* L*er* (L), *spy12* (M)*, mur1-2* (N)*, FucTA/FucTB* (O)*, spy8* (P)*, mur2-1* (R)*, mur1-1* (S)*,* 2F-Fuc treatment (U), hp*GGLT1* #1 (V) and hp*GGLT1* #3 (W)[Scale bars, 30 µm in confocal images and 2.25mm in Ruthenium red images.]

Known mutants for fucosyltransferases show no obvious adhesion defect phenotypes as opposed to *mur1-1* (S), *mur1-2* (N), treatment with 2F-Fuc (U) and hairpin silencing lines for *GGLT1* (V and W).

### Figure S2

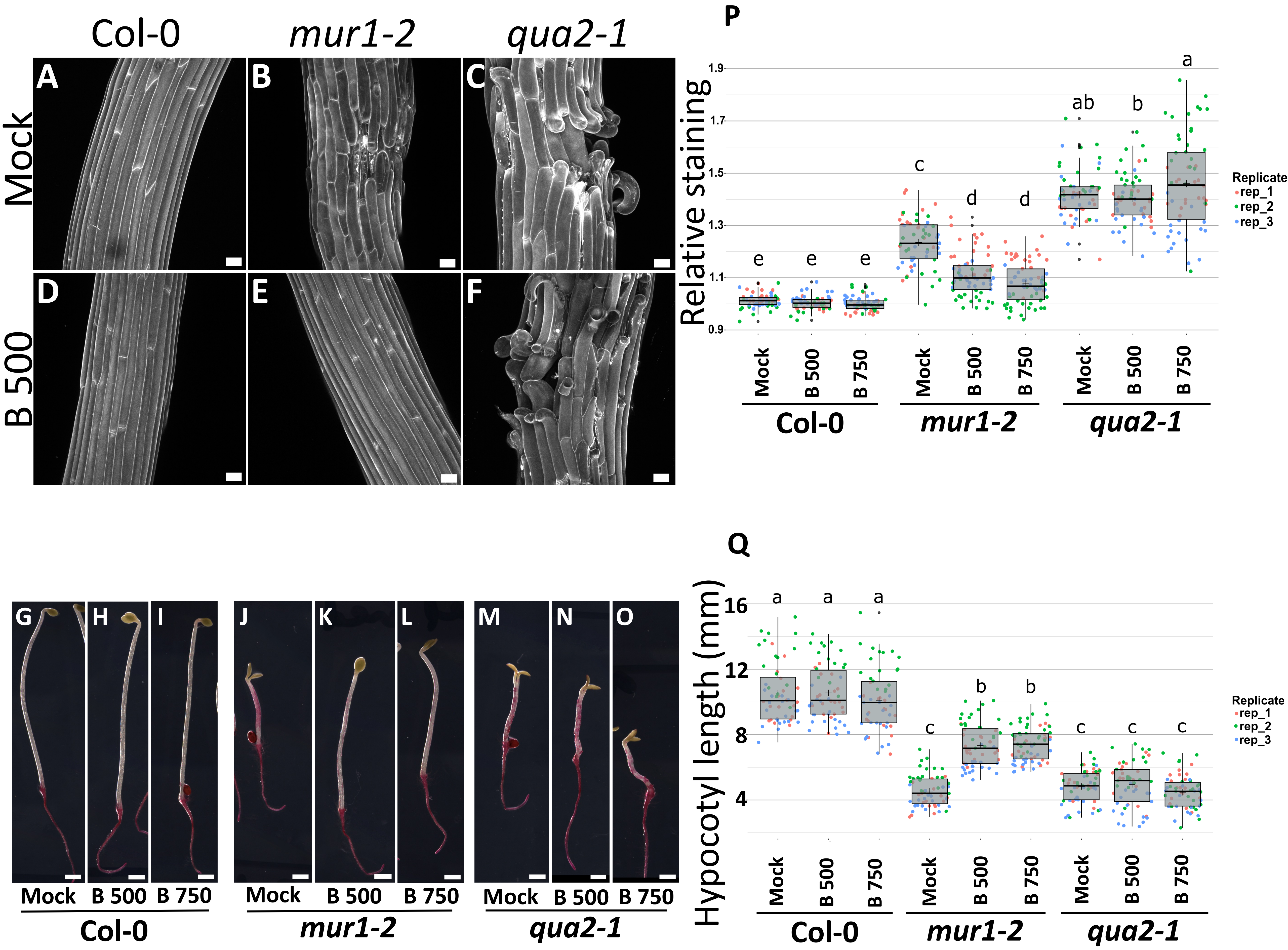


**Fig S2. Boron treatment recues cell adhesion defects in *mur1-2* but not *qua2-1*. Related to figure 1 and 2** (A-F) Representative max intensity Z-projections of confocal stacks from 4-days-old dark-grown hypocotyl stained with propidium iodide for Col-0 (A and D), *mur1-2* (B and E) and *qua2-1* (C and F) on control (A-C) and 500µM boron (D-F) containing medium. (G-O) Representative ruthenium red staining images from 4-days-old dark-grown hypocotyls from Col-0 (G-I), *mur1-2* (J-L) and *qua2-1* (M-O) on control (G,J and M), 500µM (H, K and N) and 750µM boron (I, L and O) medium. (P) Quantification of the relative ruthenium red staining intensity and (Q) measurement of hypocotyl length. Boxplots summarize three biological replicates, as represented by dots of different colors, and letters describe the statistically significant differences between populations determined by one-way ANOVA followed by Tukey's HSD test ($P < 0.05$). [Scale bars, 30 µm in confocal images and 900 µm in Ruthenium red images.]

### Figure S3

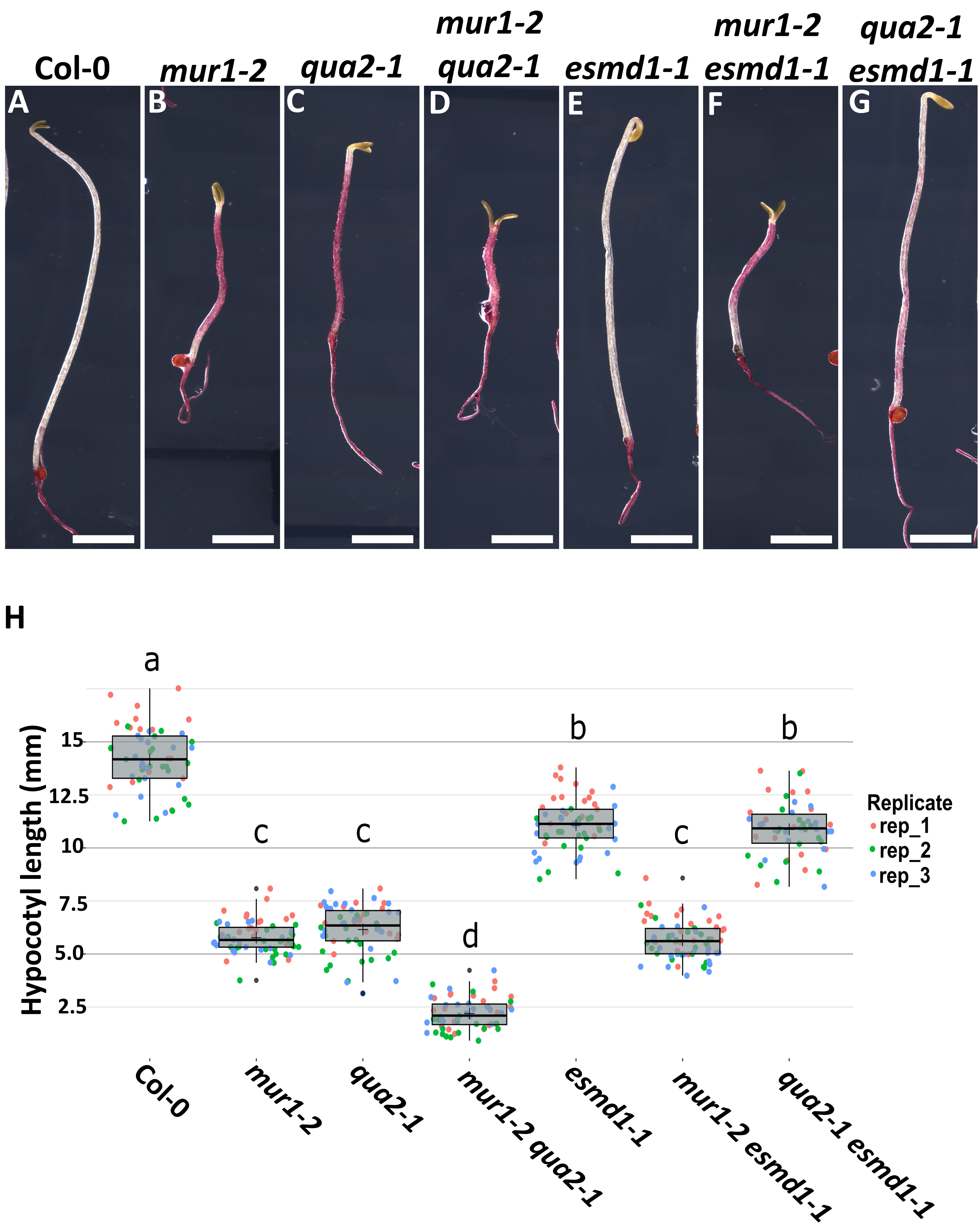


**Fig S3. Homogalacturonan deficiency and rhamnogalacturonan-II dimerization defects affect adhesion through largely separate pathways. Related to figure 2.** (A-G) Representative ruthenium red staining images from 4-days-old dark-grown hypocotyls for Col-0 (A), *mur1-2* (B), *qua2-1* (C), *mur1-2 qua2-1* (D), *esmd1-1* (E), *mur1-2 esmd1-1* (F) and *qua2-1 esmd1-1* (G). (H) Measurement of the hypocotyls length. Boxplots summarize three biological replicates, as represented by dots of different colors, and letters describe the statistically significant differences between populations determined by one-way ANOVA followed by Tukey's HSD test ($P < 0.05$).[Scale bars, 2.2mm.]

### Figure S4

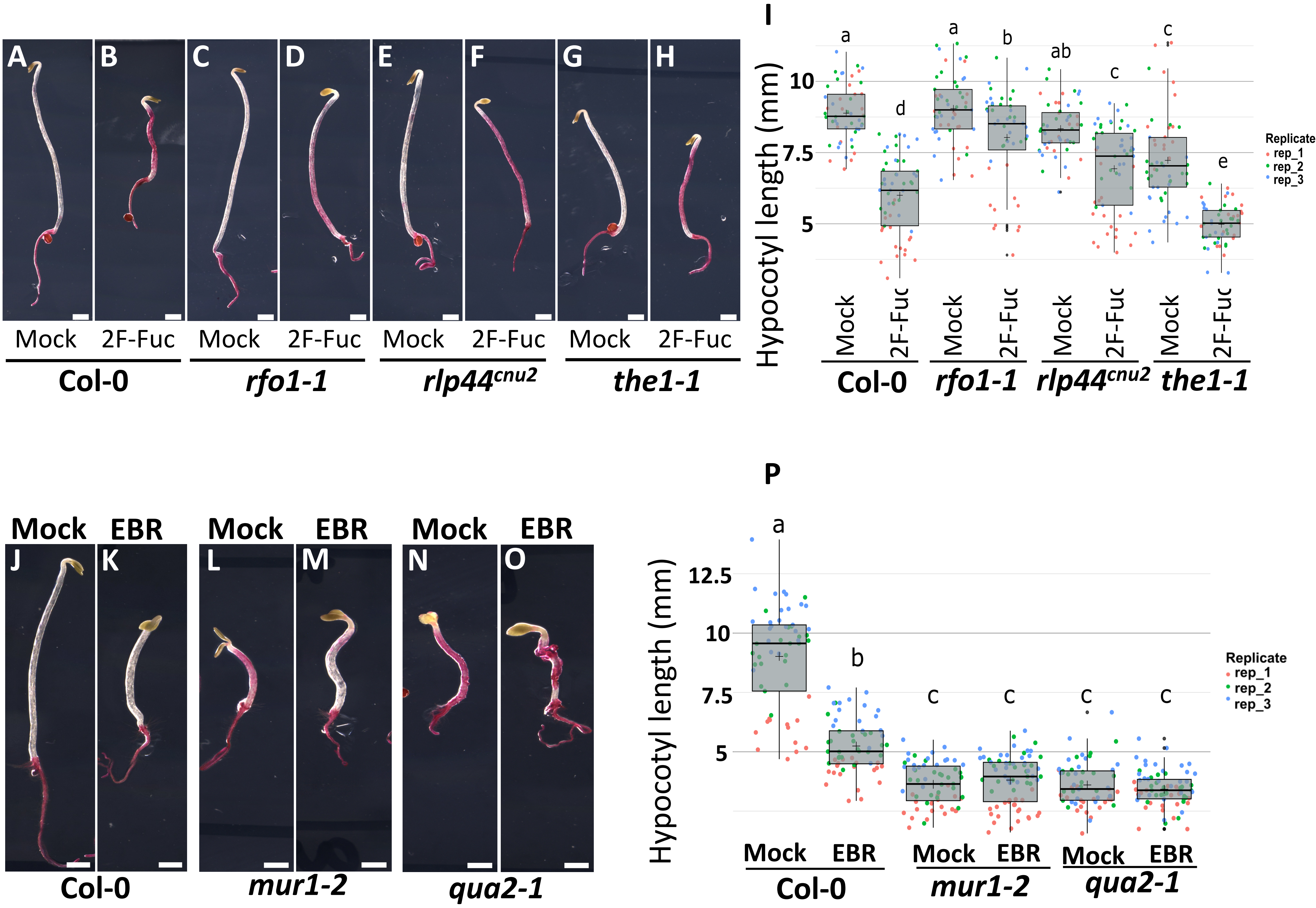


**Fig S4. Cell wall sensor mutants and brassinosteroid supplementation rescue RGII dimerization deficiency-induced adhesion defects. Related to figure 3.** (A-H and J-O) Representative ruthenium red staining images from 4-days-old dark-grown hypocotyls from Col-0 (A,B, J and K), *rfo1-1* (C and D), *rlp44*$^{cnu2}$ (E and F), *the1-1* (G and H), *mur1-2* (L and M) and *qua2-1* (N and O), under mock (A, C, E, G, J, L and N), 15µM 2F-Fuc (B, D, F and H) and 100nM EBR (K, M and O) treatment. (I and P) Measurement of the hypocotyls length. Boxplots summarize three biological replicates, as represented by dots of different colors, and letters describe the statistically significant differences between populations determined by one-way ANOVA followed by Tukey's HSD test ($P < 0.05$).[Scale bars, 900 µm.]

### Figure S5

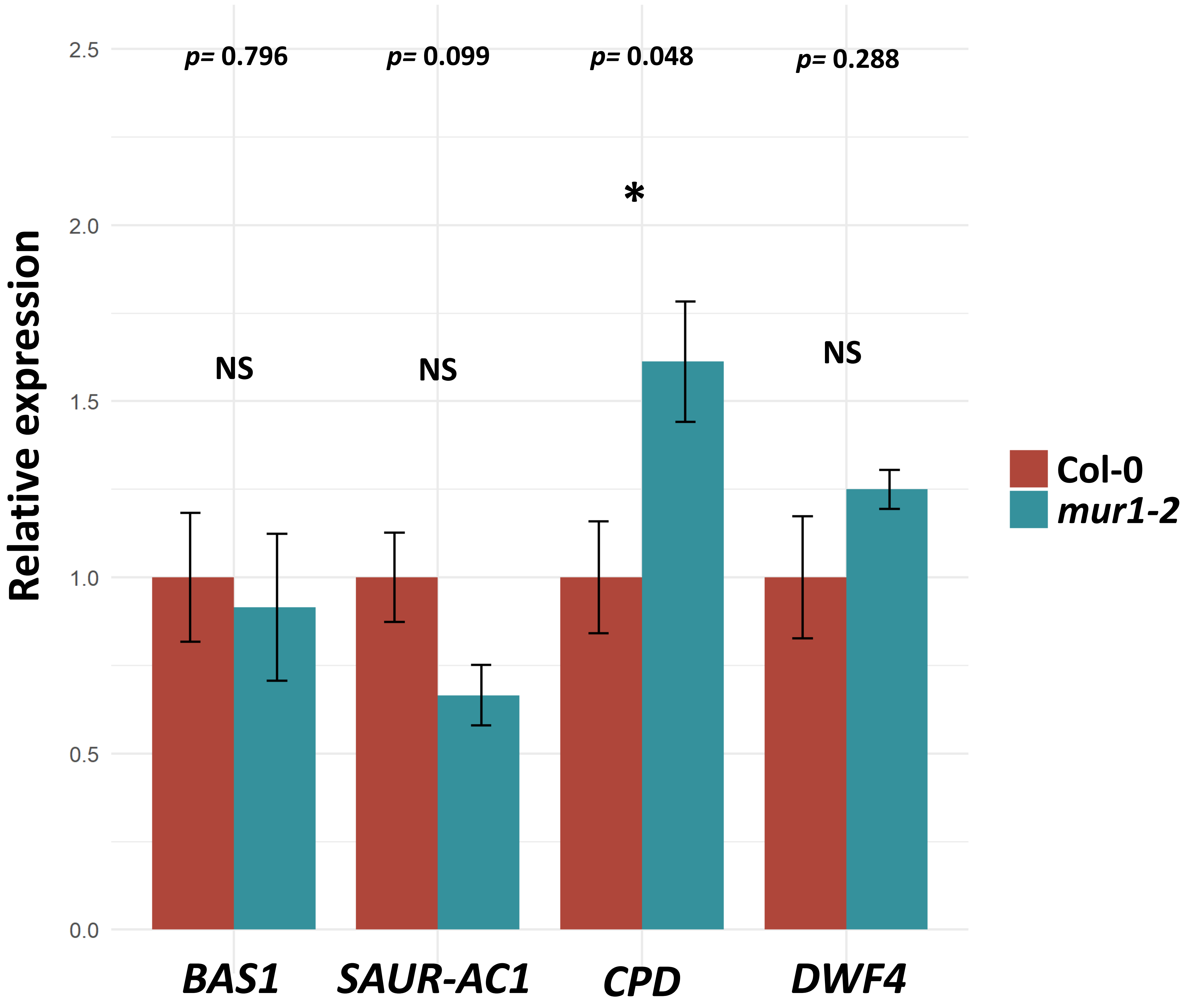


**Fig S5. Relative expression of BR responsive genes comparing Col-0 and *mur1-2.* Related to figure 3.** BAS1 and SAUR-AC1 are typically up-regulated, while CPD and DWF4 are usually down-regulated by brassinosteroid (BR) treatment. Here we observe a rather opposite pattern, suggesting a tuned-down BR pathway activity in *mur1-2*. Data are represented as mean ± SEM

### Figure S6

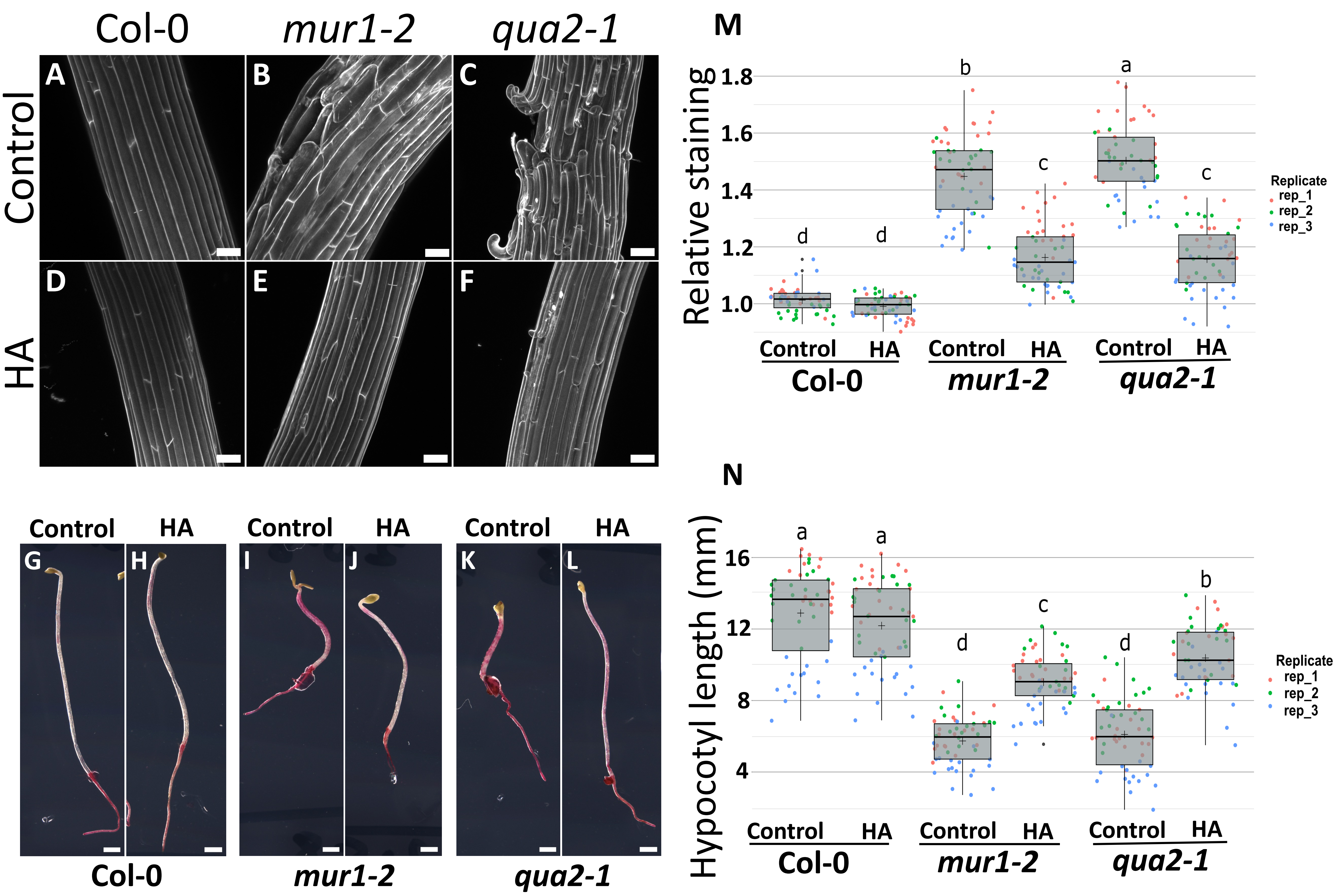


**Fig S6. Decreased water potential rescues cell adhesion defects in *mur1 and qua2-1*. Related to figure 3.** (A-F) Representative max intensity Z-projections of confocal stacks from 4-days-old dark-grown hypocotyl stained with propidium iodide for Col-0 (A and D), *mur1-2* (B and E) and *qua2-1* (C and F), on control (A-C) and high agar (HA) 2.5% (D-F) medium. (G-L) Representative ruthenium red staining images from 4-days-old dark-grown hypocotyls from Col-0 (G and H), *mur1-2* (I and J) and *qua2-1* (k and L), on control (G, I and K) and high agar (HA) 2.5% (H, J and L) medium. (M) Quantification of the relative ruthenium red staining intensity and (N) measurement of hypocotyl length. Boxplots summarize three biological replicates, as represented by dots of different colors, and letters describe the statistically significant differences between populations determined by one-way ANOVA followed by Tukey's HSD test ($P < 0.05$). [Scale bars, 30 µm in confocal images and 900 µm in Ruthenium red images.]

### Figure S7

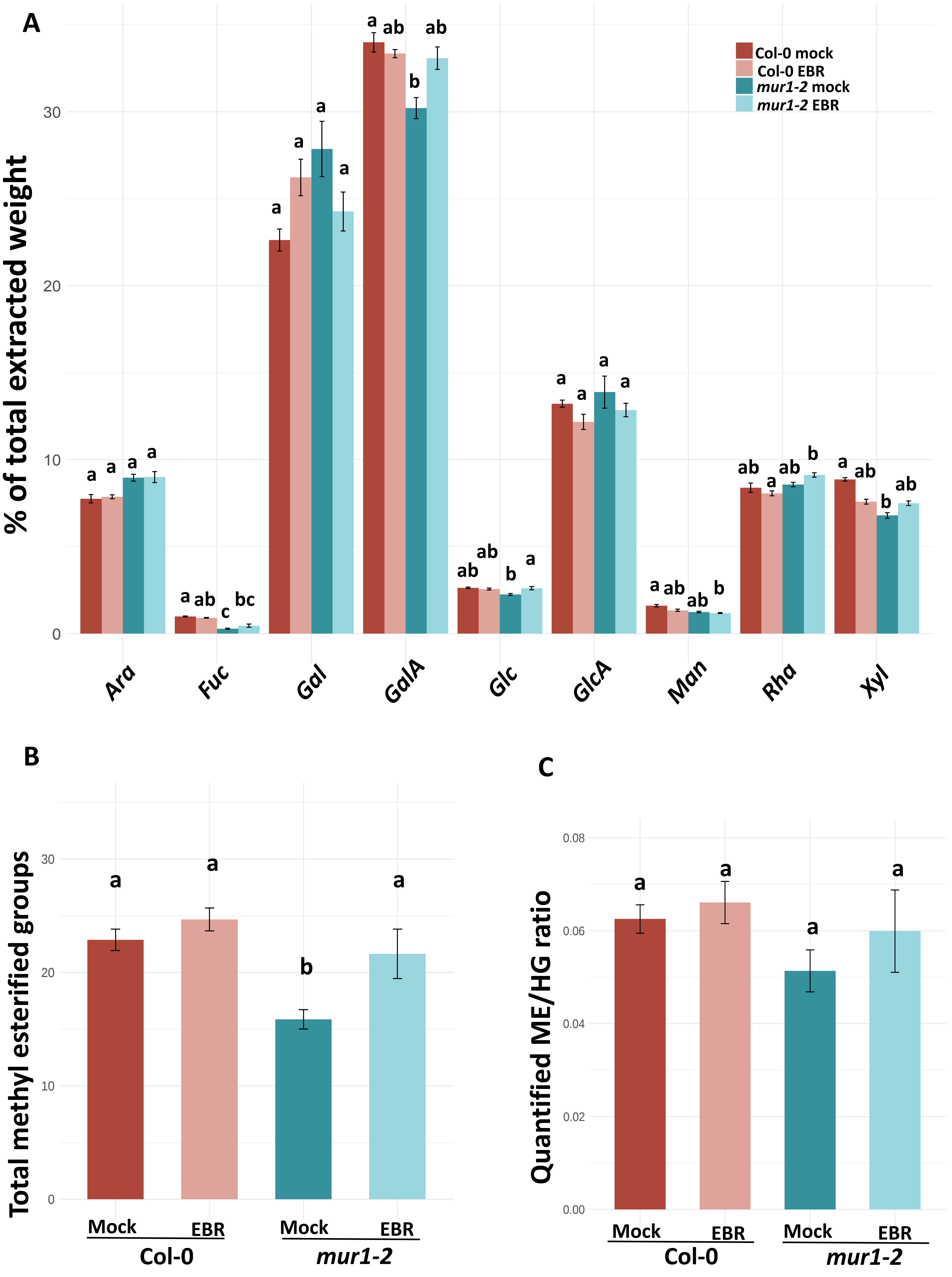


**Fig S7. Monosaccharides and methyl esters quantifications. Related to figure 4.** (A) Monosacharide content (quantity as a percentage of total extracted monosacharide weight), (B) quantified methyl esterified groups (nmol/mg dry weight) and (C) ratio of quantified methyl esterified groups over quantified homogalacturonans (See figure 4 C) for Col-0 and *mur1-2* under mock or EBR treatment. Data are represented as mean ± SEM. Letters in panels A-C describe the statistically significant differences between different populations determined in (A) by Kruskal-Wallis test followed by Dunn's test for a pairwise comparison ($P < 0.05$), (B) one-way ANOVA followed by Tukey's HSD test ($P < 0.05$) and (C) Welch's ANOVA followed by Games-Howell test ($P < 0.05$).
